# An outbreak of Rift Valley fever among peri-urban dairy cattle in northern Tanzania, 2018

**DOI:** 10.1101/2021.09.15.459147

**Authors:** William A. de Glanville, Kathryn J. Allan, James M. Nyarobi, Kate. M. Thomas, Felix Lankester, Tito Kibona, John R. Claxton, Benjamin Brennan, Ryan W. Carter, John A. Crump, Jo E.B. Halliday, Georgia Ladbury, Blandina T. Mmbaga, Furaha Mramba, Obed M. Nyasebwa, Matthew P. Rubach, Melinda K. Rostal, Paul Sanka, Emmanuel S. Swai, Agnieszka M. Szemiel, Brian J. Willett, Sarah Cleaveland

**Author notes:** **Address for correspondence:** William de Glanville, Institute of Biodiversity, Animal Health and Comparative Medicine, University of Glasgow, Glasgow, G12 8QQ, UK,; Sarah Cleaveland, Institute of Biodiversity, Animal Health and Comparative Medicine, University of Glasgow, Glasgow, G12 8QQ, UK.

## Abstract

Undetected Rift Valley fever (RVF) outbreaks are expected in endemic countries but little is known about their size or frequency. We describe a previously unreported RVF outbreak involving dairy cattle that appeared to be limited to the edge of the town of Moshi, Tanzania and occurred from May through August, 2018. The outbreak was detected retrospectively using samples collected as part of a cohort study investigating the causes of livestock abortion across northern Tanzania. A total of 14 RVF-associated cattle abortions were identified using a combination of serology and quantitative reverse transcription PCR (RT-qPCR). Milk samples from three (21%) of 14 cases were also RT-qPCR positive. Genotyping revealed circulation of RVF viruses from two lineages. The occurrence of an RVF outbreak among peri-urban dairy cattle, and evidence for RVF viral shedding in milk, highlights the potential for emerging zoonotic risks associated with the growth of urban and peri-urban livestock populations.

## Introduction

Rift Valley fever (RVF) is a mosquito-borne disease caused by Rift Valley fever virus (RVFV) that affects people and animals across Africa and the Arabian Peninsula. Previous RVF outbreaks have been major public health emergencies in affected countries and the disease is considered to be a priority for research and intervention by the World Health Organization (1). Ruminant livestock are highly susceptible to RVFV, with disease in cattle, goats, and sheep associated with abortion and mortality in young animals (2).

The epidemiology of RVF in East Africa is characterised by infrequent epidemics that are triggered by the emergence of large numbers of flood-water mosquitoes following periods of unusually heavy rainfall (3,4). In Tanzania, RVF epidemics have been reported every 10 to 15 years, with the last human or animal cases identified in the country in 2007 (5). While clinical disease in people and animals is typically not reported outside epidemics in East Africa, an increasing number of serological surveys provide evidence for regular circulation of RVFV in the region during this inter-epidemic period (IEP) (6–8).

Surveillance for RVF in most endemic countries, including Tanzania, is passive, requiring the reporting of suspicious illness in people or livestock. However, detection and confirmation of RVF cases is likely to be challenging for several reasons. In people and animals, presenting symptoms and signs of RVF are typically non-specific (2). Evidence-based thresholds for reporting and investigating unusually high rates of abortion or mortality in livestock have also not been established. Where suspect human or animal cases are reported, confirmation requires RVF-specific laboratory tests, which can have limited availability in endemic countries. These issues are compounded by medical and veterinary infrastructures being particularly weak in the grassland areas of East Africa which are typically at highest risk for RVF (3,4). Such infrastructure can be further weakened during periods of flooding which are also the periods when risk of RVFV transmission is greatest (9). Passive surveillance for RVF in these circumstances is likely to have very low sensitivity for detecting disease events (10), particularly events with small numbers of cases or with limited geographic range.

Given regular circulation of RVFV and expected low sensitivity of current surveillance for disease detection, the occurrence of small-scale outbreaks with undetected human and animal RVFV infections during periods of heavy rainfall is likely (8,9,11). Such disease events could occur regularly: in the period 2013 through 2019, meterological models predicted multiple areas of northern Tanzania to be at high risk for RVF outbreaks in all years except 2014 (12).

Here, we describe an outbreak of RVF among dairy cattle near the town of Moshi, Tanzania that occured in 2018. Infections were detected retrospectively as a result of RVF testing performed on samples collected as part of a research study investigating the infectious etiology of cattle, goat, and sheep abortions across northern Tanzania. The outbreak provides an example of the type of small-scale RVF disease event that may go undetected by national surveillance systems.

## Methods

### Prospective cohort study of livestock abortion

A prospective cohort study of livestock abortions was conducted from November, 2017 through October, 2019 in three administrative regions of northern Tanzania. This study and its principal findings are described in detail elsewhere (13). Briefly, the study population included all livestock-keeping households in 13 wards in Arusha, Manyara, and Kilimanjaro Regions. Livestock keepers in these regions have been classified as engaging in a mixture of agro-pastoral, pastoral, and smallholder-based livelihoods (14).

Each study ward was served by a livestock field officer (LFO), a government-employed veterinary technician providing basic veterinary services. Community meetings were held in study wards and livestock keepers were asked to report any cattle, goat, and/or sheep abortions to their LFO. Within 72 hours of a reported abortion, the LFO collected blood, milk, and vaginal swab samples from the aborting dam. Placental tissue samples and fetal surface swabs were also collected where available. Tissue, milk, and swab samples were preserved in DNA/RNA shield (Zymo Research, California, USA). Basic individual animal- and household-level data, including breed, service date, animal demographics, and vaccination history were recorded.

All households were re-visited four to six weeks after the investigated abortion and a convalescent-phase blood sample collected from the affected dam.

### RVF livestock abortion case definition

Based on World Organisation for Animal Health (OIE) guidelines (15), we used a combination of molecular and serological methods to identify RVF cases. A case of RVF-associated livestock abortion was defined as an abortion event with RVFV detected by one-step reverse-transcription real-time quantitative PCR (RT-qPCR) in any of the vaginal swab, fetal swab, or placental tissues, and evidence of RVFV antibodies by ELISA on a blood sample collected on an acute or convaelescent serum sample from the dam (13).

RVF molecular and serological testing was performed retrospectively, with diagnoses made approximately 12 months after abortion events occurred. Households in which RVF cases were identified were revisited shortly after detection (i.e., around 12 months post abortion) and livestock keepers informed of positive test results. During this third visit, a blood sample was collected from all animals that had been confirmed as RVF cases and which were still present.

### Serological testing for RVFV

Sera were separated from clotted whole blot samples by centrifugation (1,300 x g for 10 mins). Sera from acute, convalescent and 12-month blood samples were tested for RVFV IgG antibodies using a multi-species competitive ELISA (ID Screen, IDVet, Paris, France).

### Reverse transcriptase quantitative polymerase chain reaction

RNA was extracted from swab, tissue, and milk samples using a RNeasy® Mini kit (QIAGEN, Hilden, Germany) as described by Thomas et al 2021 (13). Detail on extraction from milk samples is given in the Supplementary Materials. Detection of RVFV by RT-qPCR on extracted RNA was performed using published protocols (16). Briefly, a 94 base-pair target of the RVFV G2 gene was amplified using the primer pair RVS and RVA and a fluorescent-labelled probe (RVP). A negative extraction control, negative PCR control (PCR grade water) and positive control (RNA extracted from cells experimentally infected with RVFV-MP12 (17)) were included in each PCR run. Samples were run in duplicate with crossing threshold (Ct) values <40 on both wells considered positive.

### Genotyping of RVFV

RNA from positive RT-qPCR samples was shipped to the University of Glasgow for viral genotyping. An approximately 900bp target of the S-genome segment was amplified using previously described primers (18). Reactions were performed using the Qiagen® OneStep RT-PCR kit using 8 μl of extracted RNA in a final reaction volume of 25 μl. Cycling conditions were reverse transcription at 50°C for 60-90 minutes followed by an initial PCR activation step of 95°C for 15 minutes and then 45 cycles of 94°C for 1 minute, 60°C for 1 minute and 72°C for 1.5 minutes. A final extension step of 72°C for 10 minutes was performed. A negative control (PCR grade water) and positive control (RVFV-MP12) were included in each run. PCR products were visualised by gel electrophoresis. For sequencing, PCR products from positive samples were purified using the QIAquick™ Gel Extraction kit. Sanger sequencing was performed using both forward and reverse primers by SourceBioscience (Nottingham, UK) using dGTP technology.

Molecular phylogenetic analysis was performed in MEGA7.0 (19). Sequences were aligned using the Clustal W algorithm (19). The evolutionary history was inferred using the maximum likelihood method based on the Kimura 2-parameter model with a discrete Gamma distribution (20). RVFV sequences were compared to sequences from a diverse set of RVFV strains through BLAST searches of the GenBank nucleotide database to identify the infecting haplotype (21). Particular attention was paid to viral sequences that fell into the same haplotype clade as the live vaccine used in the region (Smithburn RVFV modified vaccine strain, GenBank Accession number DQ380157) as these can induce abortions in pregnant animals (22). The tree with the highest log likelihood value was selected and drawn to scale with branch lengths measured in the number of substitutions per site. All positions containing gaps or missing data were deleted.

### Research ethics

The study protocols, questionnaires, and consent documents were approved by the Kilimanjaro Christian Medical Centre (KCMC) (832) and National Institute of Medical Research (NIMR) (2028) research ethics committees, the University of Otago Ethics Committee (H17/069), and University of Glasgow Medical, Veterinary and Life Sciences (MVLS) ethics committee (200140152 and 20017006). Permission to carry out the study in Tanzania was provided by the Tanzania Commission for Science and Technology (2014-244-ER-2005-141). Publication of this work has been approved by the Director of Veterinary Services, Tanzania.

## Results

### Prospective cohort study of livestock abortion

Samples were collected from 215 abortion events from November, 2017 through October, 2019. Of the investigated abortions, 71 (33%) involved cattle, 100 (47%) goats, and 44 (21%) sheep. Additional detail on breed of affected animals is given in the Supplementary Materials. Both RT-qPCR and acute or convalescent phase serological testing for RVF were performed on samples from 212 (99%) of 215 abortion events. The number of abortion events investigated per ward was highly variable, with a minimum of one and maximum of 84 abortion events investigated per ward (Table S1, Supplementary Information).

### Detection of RVF livestock abortion cases

In total, 14 abortion events met our case definition for RVF-associated abortion. All occurred in cattle between 16^th^ May and 11^th^ August 2018. These 14 cases were found in 14 households and represented 23% of all 63 abortion investigations conducted as part of the prospective cohort study over this time period (i.e., the apparent outbreak period) (Figure 1). Case animals were all found in four wards in three districts surrounding the town of Moshi in Kilimanjaro Region (Figure 2). These were Arusha Chini and Kindi Wards in Moshi Rural District; Rau Ward in Moshi Municipality; and Machame Mashariki Ward in Hai District.

**Figure 1.**
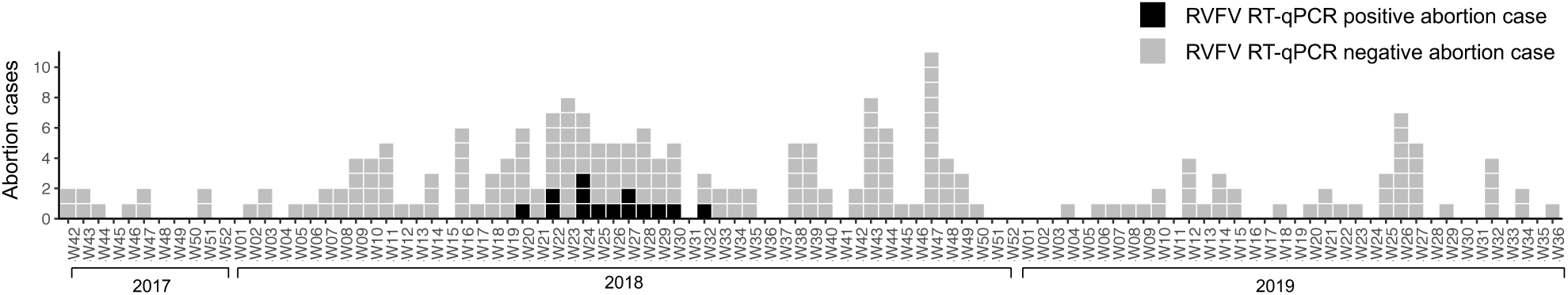
RVFV RT-qPCR status of livestock abortions in northern Tanzania by week, November 2017 to October 2019.

**Figure 2.**
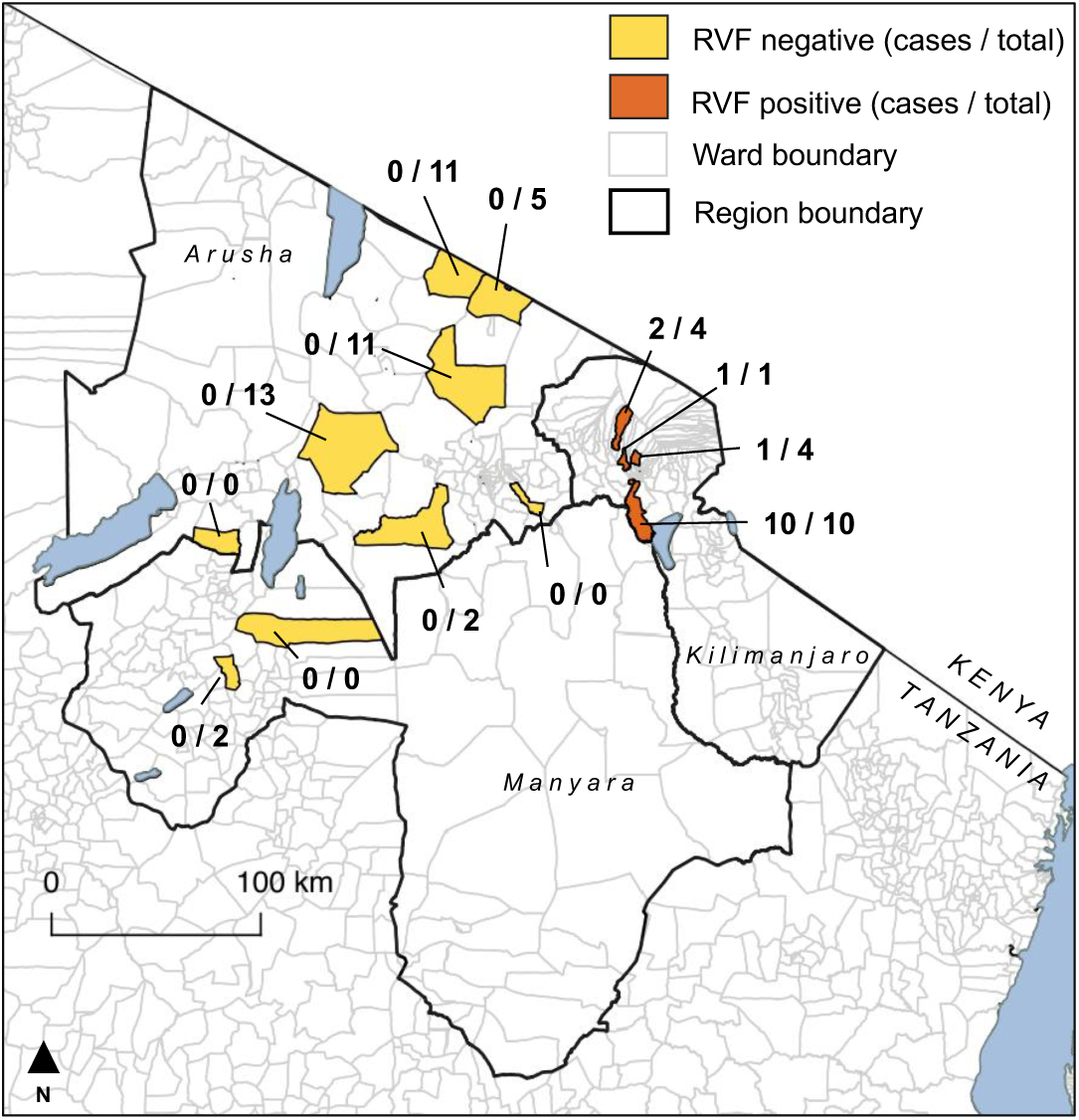
Location of wards included in the prospective cohort study of livestock abortion in northern Tanzania, 2017-2019. The number of RVF cases identified out of total abortions investigated during the apparent outbreak period (16^th^ May through 11^th^ August, 2018) are shown.

Table 1 shows the samples collected from case animals and the results of RT-qPCR and serological testing performed. Further detail on crossover thresholds for RT-qPCR and percentage competition ELISA values are given in the Suppementary Materials (Table S2 and S3). Milk samples and vaginal swabs were available from all 14 cases, of which three (21%) and 11 (79%) were positive by RT-qPCR, respectively. Ten (91%) of 11 fetal swabs and all six (100%) placental samples from confirmed cases were positive for RVFV by RT-qPCR. Of 14 RVF abortion cases, 13 (93%) dams were seropositive on the basis of samples collected within 72 hours of abortion. The one (7%) further case demonstrated seroconversion between acute and convalescent sera (Table 1). All seven case animals that could be sampled 12-months post abortion remained seropositive (Table 1 and Table S2).

**Table 1.** Results of RT-qPCR and serological testing performed on samples from confirmed cattle RVF cases in northern Tanzania May through July 2018. + positive sample; - negative sample; NA = sample type that was not available; * samples used for genotyping (individual identifiers: ^1^SEBI-051; ^2^SEBI-079; ^3^SEBI-094)

| Abortion<br>date (2018) | District | RT-qPCR |  |  |  | Serology |  |  |
| --- | --- | --- | --- | --- | --- | --- | --- | --- |
|  |  | Milk | Vaginal<br>swab | Placental<br>sample | Fetal<br>swab | Acute<br>serum | Convalescent<br>serum | 12-month<br>serum |
| 16 May | Moshi Rural | - | + | NA | +*<sup>1</sup> | - | + | + |
| 29 May | Moshi Rural | + | + | NA | + | + | + | + |
| 29 May | Moshi Rural | + | + | NA | + | + | + | + |
| 12 Jun | Moshi Rural | - | + | NA | + | + | + | + |
| 11 Jun | Moshi Rural | - | + | + | + | + | NA | + |
| 17 Jun | Moshi Rural | + | - | +*<sup>2</sup> | - | + | + | NA |
| 24 Jun | Moshi Rural | - | - | + | + | + | + | NA |
| 29 Jun | Moshi Rural | - | + | NA | NA | + | + | + |
| 03 Jul | Moshi<br>Municipality | - | + | NA | +*<sup>3</sup> | + | + | + |
| 03 Jul | Moshi Rural | - | + | NA | NA | + | NA | + |
| 15 Jul | Hai | - | - | + | NA | + | + | NA |
| 22 Jul | Moshi Rural | - | + | NA | + | + | + | + |
| 26 Jul | Hai | - | + | + | + | + | + | + |
| 11 Aug | Moshi Rural | - | + | + | NA | + | + | NA |

One cow that aborted in August 2018 in Hai District and one cow that aborted in July 2019 in Moshi Rural District were also seropositive for RVFV on acute serum samples (and the one available convalescent sample from these animals). Both animals were negative by RT-qPCR testing of milk, vaginal swabs, and fetal swabs and so did not meet the case definition for RVF-associated abortion.

### Characteristics of RVF cases and affected households

Of the 14 RVF case cattle, one (7%) was an indigenous breed, seven (50%) were European dairy breeds and six (43%) were cross-breeds. Service date was known for 12 case animals, with abortions occurring at a median (range) of 181 (128 to 251) days of gestation. No RVF vaccination was reported in any of the affected herds in the previous 24 months. The median (range) number of adult female cattle kept by households with RVF cases was five (1 to 14). Seven of the affected households reported keeping goats in addition to cattle, with a median of one (0 to 30) adult female goats kept. No other livestock were kept by affected households.

Data on cattle grazing management were available for 12 (71%) of the 14 households with RVF cases. Of the households with these data, five (42%) managed cattle under a zero-grazing system (permanently housed with fodder cut and brought to animals), three (25%) managed cattle under a tethered grazing system (unhoused but restrained by long rope and moved between grazing areas near to households), and four (33%) managed cattle under a herded grazing system (unhoused and moved between grazing areas by a herdsperson). One (8%) household reported using a mix of zero- and herded-grazing systems for cattle.

### RVFV genotypes identified

An 802-bp fragment of RVFV genome Segment S sequence was obtained from three RVF cases. Positive cattle came from Moshi Rural (SEBI-051 and SEBI-079) and neighbouring Moshi Municipality (SEBI-094) Districts.

Sequences obtained from SEBI-079 and SEBI-094 were highly similar (99.9%) with only a single nucleotide substitution between the two sequences. Based on phylogenetic analysis, these two sequences fell within the Clade B haplotype (21) alongside numerous sequences obtained from previous RVF outbreaks in livestock in Kenya, Tanzania, and Sudan (Figure 3). In contrast, the third sequence (SEBI-051) fell within the Clade E haplotype and was more similar to sequences obtained from a South African outbreak in 2009 than sequences previously described in East Africa (Figure 3). Despite falling into the same clade as the Smithburn vaccine strain, the predicted protein encoded by this sequence did not contain any of the amino-acid substitutions associated with the modified vaccine strain and was therefore considered to be a wild-type virus.

**Figure 3.**
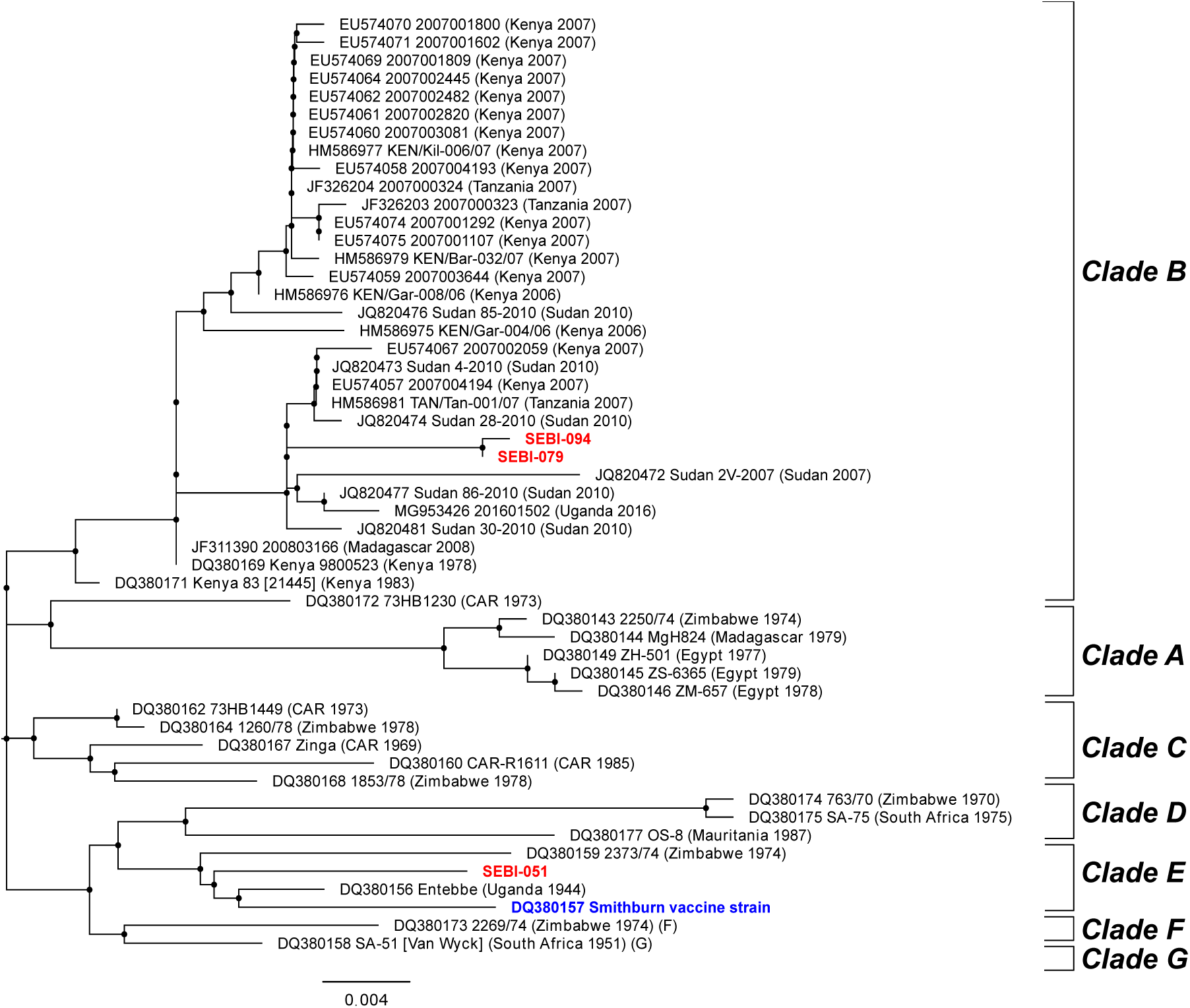
Molecular phylogenetic analysis of partial sequence (802bp) of Segment S of Rift Valley fever virus strains obtained from Tanzanian cattle, northern Tanzania, 2018. *The tree with the highest log likelihood (−2414*.*72) is shown. Sequences are labelled with Genbank accession number and designated strain names with country and year of origin shown in round parentheses. Sequences from this study are highlighted in red and labelled with Genbank accession numbers (pending) and unique animal individual identifiers. Clades of viral lineages as designated by Bird et al (21) are labelled by square parentheses. Sequence from the Smithburn vaccine strain is highlighted in blue*.

## Discussion

We describe an apparently small outbreak of RVF occurring among dairy cattle in a peri-urban area of northern Tanzania in 2018. Detection of livestock cases was accomplished through active, abortion-based sampling triggered by individual animal abortions rather than through a passive surveillance response to reports of unusually high levels of abortion. The retrospective detection of this apparently small, previously unreported RVF outbreak using samples collected as part of a research study provides evidence that RVF-associated livestock abortions are occurring more regularly in this endemic setting than would be expected on the basis of national surveillance alone. The occurrence of small-scale RVF outbreaks that go undetected by surveillance systems in endemic settings is not unexpected but such events have rarely been described in detail.

The outbreak we describe has several notable features. First, all RVF cases were detected in animals reared in a peri-urban area. Transmission in urban and peri-urban areas is of key public health concern for many arboviruses (23) but has been infrequently described for RVFV. Indeed, the World Health Organization reports that there is no evidence for RVF outbreaks in urban areas (24), which could be taken to imply that RVF does not pose a threat to urban populations. In Tanzania, the peri-urban livestock sector is undergoing rapid expansion to meet growing demand for milk and meat (25). Growing livestock populations within and surrounding high human population density areas may pose emerging risks for zoonotic disease transmission (26), including for for RVF.

These zoonotic risks are highlighted by the detection of RVFV nucleic acids in milk from multiple aborting dams sampled in this study. All RVF-associated abortions we identified were dairy cattle reared in an area that supplies the majority of milk to the town of Moshi (27). To our knowledge, the only published report of RVFV detection in milk was from experimentally infected animals in the 1950s (28). Our samples were preserved in a viricidal solution so we were unable to assess the infectivity of milk from RVF cases, but milk-borne RVFV transmission is supported by literature-based reports of significant positive associations between RVFV seropositivity in people and a history of un-boiled milk consumption (29). Few, if any, data exist on the effectiveness of heat-treatment for inactivation of RVFV in milk and milk products. RVFV viability using culture-adapted virus was detected following short periods of heat treatment at 56 °C (30), raising concerns about the potential thermal stability of RVFV. In heat treatment experiments performed by our group using the RVFV-MP12 vaccine strain spiked in milk, we found no viral survival after incubation at 72 °C for 15 minutes (further detail in Supplementary Materials). Other authors report complete inactivation of RVFV after heating solutions containing the virus for one hour at 60 °C (31). These data suggest heat treatment is likely to reduce risks of RVFV transmission via milk but further work is needed to explore the effectiveness of standard milk pasteurisation and other milk preparation procedures on inactivation of the virus. Milk can be an important source of nutrition for food insecure households in RVF endemic settings and interventions aiming to promote milk safety should be carefully designed to avoid unintended consequences such as reduced consumption or inequitable access to safe products (27).

A second notable feature of this outbreak is that despite its apparently limited spatial and temporal extent, we identified circulation of viruses from two distinct lineages. The outbreak around Moshi occurred at the same time as outbreaks involving human and livestock populations were occurring in Kenya, Rwanda, and Uganda (10,32). It is therefore possible that there were repeated virus introductions via livestock movments from neighbouring countries, which could explain the co-occurrence of these two viral haplotypes. This would also suggest that the Moshi outbreak was part of more widespread, regional-level RVFV transmission rather than a small, isolated RVF outbreak arising autochthonously in northern Tanzania. Alternatively, the apparent viral diversity observed could be explained by repeated emergence from the low-level endemic viral cycling among livestock that has been described in Tanzania (6–8) or spillover from unknown sylvatic cycles. It is worth noting that the town of Moshi sits on the edge of Kilimanjaro National Park, and one of the wards in which cases were identified (Machame Mashiriki) directly borders this forested area. While limited data exist, it has been suggested that RVFV may circulate in unidentified sylvatic cycles in the forests of East Africa (33). Attempts to confirm and characterise these cycles in northern Tanzania would represent a valuable area for future research. At the time of writing, we are not aware of viral sequence data from the outbreaks in the wider East African region in 2018 that could help address questions around virus origin. These unanswered questions, and the diversity of viral strains we observed, clearly demonstrate the value of genotyping during RVF oubreaks (34).

While there are many logistic and economic challenges to collecting diagnostic material from animal abortions in low-resource settings, we show that molecular evidence for RVFV infection can be detected in a range of sample types collected within 72 hours of abortion. The number of available cases was small, and all sample types were not available for all animals, but placental material was most consistently RT-qPCR positive among the RVF cases, followed by swabs from the fetal surface. Encouragingly, vaginal swabs from aborted dams appear to have good sensitivity for RVFV detection. Sampling directly from the dam has considerable benefits over collecting swabs or material from the fetus or placenta, which may be scavenged by dogs and wildlife or may become rapidly autolysed in hot climates (35).

The small number of cases detected in this study, and the apparent geographic isolation of cases, suggests the occurrence of a small-scale RVF outbreak. However, there are important limitations to the prospective cohort study in which RVF cases were detected that create uncertainty about the true size of the outbreak we describe. In particular, there were substantial differences in the number of abortions investigated in each study ward, with reporting likely to be strongly biased by levels of engagement of community members with their LFO (13). A particular source of this bias that is reflected in our data is the large numbers of European dairy breeds and their crosses in the study sample and among cases. These animals are typically higher value than local breeds and can therefore be expected to receive higher levels of veterinary input. European-breed dairy cattle and their crosses are primarily reared in smallholder, peri-urban areas in northern Tanzania and are relatively rare across rural areas of northern Tanzania (14). Smallholder dairy systems should therefore be considered to be over-represented in our sample. Given this selection bias in our sample, we cannot rule out a larger RVF outbreak or multiple separate outbreaks occurring over a wider geographic area or longer time period. All RVF abortions were detected during a high rainfall period in which multiple areas of northern Tanzania were considered to be at high risk for RVF outbreaks (12,32).

## Conclusion

Testing of samples collected as part of a research study investigating the etiology of livestock abortion allowed us to retrospectively detect an outbreak of RVF occurring among livestock in Tanzania in 2018. Our data provide a rare example of the type of small-scale RVF outbreak that can occur below the surveillance detection threshold in endemic countries. We identified shedding of RVFV nucleic acids in milk from affected animals, supporting growing evidence for milk as a potential source of human RVFV infection. Promotion of milk safety measures, including pastuerisation and home boiling, could be expected to contribute to reduced zoonotic RVF risk in endemic settings during outbreaks. The likely occurrence of undetected outbreaks during high risk periods suggest that the promotion of such measures should include high risk periods, in addition to during known outbreaks. Promotion of milk safety measures to reduce RVF risk should also include urban supply chains.

## Acknowledgements

We would like to thank participating livestock-keepers, as well as village, ward, district and regional authorities. Our particular thanks to the LFOs who collected data for this study. We are also very grateful to Rigobert Tarimo, Fadhili Mshana, Matayo Melubo, Hassan Hussein, Ephrasia Hugho, Nelson Amani, Elizabeth Kasagama, Victor Mosha and Rosanne de Jong for their contribution to field and/or laboratory work. This research was supported by the Supporting Evidence Based Interventions project, University of Edinburgh (R83537) and the Zoonoses and Emerging Livestock Systems program (funded through BBSRC, DfID, ESRC, MRC, NERC and DSTL) (BB/L018926/1, BB/L017679/1, BB/N503563/1). Additional financial support was provided through BBSRC (BB/J010367) and the National Institutes of Health (R01TW009237).

## Notes

### Competing Interest Statement

The authors have declared no competing interest.

