## Supplementary Materials for "An outbreak of Rift Valley fever among peri-urban dairy cattle in northern Tanzania, 2018"

**Corresponding authors:**

### Additional detail on RNA extraction from milk

Milk samples were preserved in equal volumes of double concentration DNA/RNA shield. A 200µl sample with 600µl RLT buffer and 10µl Proteinase K was heat treated at 55°C for 10 minutes. When cool, 300µl ethanol (96%) was added to the tube and the RNA from this tube extracted using the RNeasy Mini kit (QIAGEN, Hilden, Germany) following manufacturer's instructions.

### Additional detail on the prospective cohort study sample

The number of abortion investigations investigated in study wards as part of the prospective cohort study is shown in Table S1. Also shown is the number of abortion investigations performed over the period between the first (16th May 2018) and the last (11th August 2018) RVF positive case and the number of RVF cases identified in each ward.

The prospective cohort study collected diagnostic samples from 215 individual animal abortion events, of which 71 (33.0%) were cattle, 100 (46.5%) were goats, and 44 (20.5%) were sheep. Of the cattle, 16 (22.9%) were indigenous breeds, 17 (24.9%) were European (exotic) dairy breeds, and 36 (51.4%) were crosses between the two. One bovine animal was an unclassified breed. Of the goats, 77 (77.0%) were indigenous breeds, 3 (3.0%) exotic breeds, and 17 (17.0%) were crosses. Three were unclassified. Of the sheep, 41 (91.1%) were indigenous breeds and 3 (6.7%) were crosses. One was unclassified.

**Table S1.** Number of RVF cases identified and abortion investigations conducted in each ward included in the prospective cohort study in northern Tanzania between November 2017 through October 2019.

| Ward | District | RVF cases | Outbreak period total investigations[a] | Study period total investigations[b] |
|---|---|---|---|---|
| 1. Arri | Babati | 0 | 2 | 2 |
| 2. Arusha Chini | Moshi Rural | 10 | 10 | 14 |
| 3. Engarenaibor | Longido | 0 | 11 | 23 |
| 4. Engikaret | Longido | 0 | 11 | 28 |
| 5. Kansay | Karatu | 0 | 0 | 1 |
| 6. Kikwe | Meru | 0 | 0 | 2 |
| 7. Kimokouwa | Longido | 0 | 5 | 5 |
| 8. Kindi | Moshi Rural | 1 | 1 | 1 |
| 9. Machame Mashariki | Hai | 2 | 4 | 37 |
| 10. Magugu | Babati | 0 | 0 | 1 |
| 11. Meserani | Monduli | 0 | 2 | 5 |
| 12. Selela | Monduli | 0 | 13 | 84 |
| 13. Rau | Moshi Municipality | 1 | 4 | 12 |
| Total | | 14 | 63 | 215 |

[a] Total number of animals sampled during the period between the first and last confirmed RVF case; [b] Total number of animals sampled over the whole study period.

**Additional detail on results on diagnostic testing performed on RVF cases**

A breakdown of the ELISA percentage competition values is given in Table S2. All animals except one were seropositive during the first investigation (i.e. within 72 hours of the abortion). It was possible to collect 12-month follow-up serum samples from 9 RVF cases of which all were seropositive with minimal change in percentage competition values when compared to acute and/or convalescent samples. A breakdown of the RT-qPCR crossover threshold (Ct) values for samples from RVFV positive animals identified in the prospective cohort study is given in Table S3.

**Table S2.** Values for ELISA percentage competition for serum samples collected from confirmed RVF cases in northern Tanzania in 2018. Samples with a percentage competition <40 are considered seropositive.

| ID | Acute serum | Convalescent serum | 12-month serum |
|---|---|---|---|
| SEBI-051 | 98.3* | 3.9 | 4.1 |
| SEBI-058 | 9 | 3.5 | 3.9 |
| SEBI-064 | 8.4 | 4.5 | 3.9 |
| SEBI-075 | 6.6 | 3.6 | 3.8 |
| SEBI-076 | 4.0 | NA | 33.6 |
| SEBI-079 | 12.0 | 3.4 | NA |
| SEBI-084 | 7.1 | 3.4 | NA |
| SEBI-089 | 5.3 | 4.0 | 3.8 |
| SEBI-094 | 6.4 | 3.7 | 4 |
| SEBI-095 | 7.8 | NA | 4.5 |
| SEBI-099 | 5.0 | 3.6 | NA |
| SEBI-106 | 3.8 | 3.2 | 5.0 |

| | | | |
|---|---|---|---|
| SEBI-110 | 5.8 | 3.9 | 4.1 |
| SEBI-115 | 5.5 | 3.4 | NA |

*Seronegative; NA sample not collected

**Table S3.** Crossover threshold (Ct) values from paired RVFV RT-PCR positive samples from RVF-associated abortion cases occurring in northern Tanzania, May through August 2018.

| ID | Milk Ct 1 | Milk Ct 2 | Vaginal swab Ct 1 | Vaginal swab Ct 2 | Placenta Ct 1 | Placenta Ct 2 | Fetus swab Ct 1 | Fetal swab Ct 2 |
|---|---|---|---|---|---|---|---|---|
| SEBI-051 | - | - | 31.94 | 31.92 | *NA* | *NA* | 24.74 | 23.97 |
| SEBI-058 | 34.39 | 34.20 | 39.0 | 38.74 | *NA* | *NA* | 39.26 | 39.42 |
| SEBI-064 | 35.16 | 34.62 | 28.75 | 28.68 | *NA* | *NA* | 26.47 | 25.94 |
| SEBI-075 | - | - | 33.54 | 33.89 | *NA* | *NA* | 18.37 | 18.43 |
| SEBI-076 | - | - | 39.76 | 39.03 | 26.04 | 26.04 | 23.25 | 22.93 |
| SEBI-079 | 30.82 | 30.41 | - | - | 27.47 | 27.31 | - | - |
| SEBI-084 | - | - | - | - | 22.26 | 22.42 | 21.68 | 21.68 |
| SEBI-089 | - | - | 31.11 | 31.04 | *NA* | *NA* | *NA* | *NA* |
| SEBI-094 | - | - | 28.56 | 28.97 | *NA* | *NA* | 19.09 | 19.04 |
| SEBI-095 | - | - | 31.0 | 31.9 | *NA* | *NA* | *NA* | *NA* |
| SEBI-099 | - | - | - | - | 18.11 | 18.15 | *NA* | *NA* |
| SEBI-106 | - | - | 30.6 | 30.62 | *NA* | *NA* | 21.72 | 21.77 |
| SEBI-110 | - | - | 27.22 | 27.07 | 24.98 | 25.08 | 23.23 | 23.42 |
| SEBI-115 | - | - | 31.0 | 31.28 | 21.29 | 21.63 | *NA* | *NA* |

NA = samples that were not available; '-' are RT-PCR negative results.

**Heat treatment experiment**

We tested survival of recombinant RVFV MP12 strain in milk following heat-treatment. Virus stocks were grown at 37 °C in Vero-E6 cells in Dulbecco's modified Eagle's medium (DMEM) supplemented with 2 % foetal calf serum (FCS) by infecting at a multiplicity of infection (MOI) of 0.001 and harvesting the culture medium when cytopathic effect was observed. Virus stock (2 x $10^7$ pfu/ml) dilutions (1/10, 1/100, 1/1000, 1/10000) were prepared in 1 ml of raw milk sourced from cattle in the UK in two triplicate batches. One batch was placed in a heat block set to 72 °C. After the samples reached the temperature of 72 °C (approximately 6 minutes), they were incubated for 15 minutes. Samples were then allowed to cool to room temperature. The second batch was kept at room temperature as negative control. To assess RVFV viability all the samples were titrated by plaque assay. Briefly, Vero E6 cells in 12 well plates were infected with serial dilutions of samples made in DMEM containing 2 % FCS. After 1 h at 37 °C, inoculum was removed and overlay comprising DMEM supplemented with 2 % FCS, 1x Antibiotic Antimycotic Solution (Sigma, A5955) and 0.6 % Avicell (FMC) was added. Plates were incubated at 37 °C for 3 days. Cell monolayers were fixed with 8 % formaldehyde and plaques were visualised by staining with crystal violet. All experiments with infectious RVFV MP12 strain were conducted under biosafety level 3 (BSL-3) conditions at the University of Glasgow.

No viral plaques were detected following the heat treatment of milk containing RVFV MP12 strain at multiple dilutions (Figure S1).

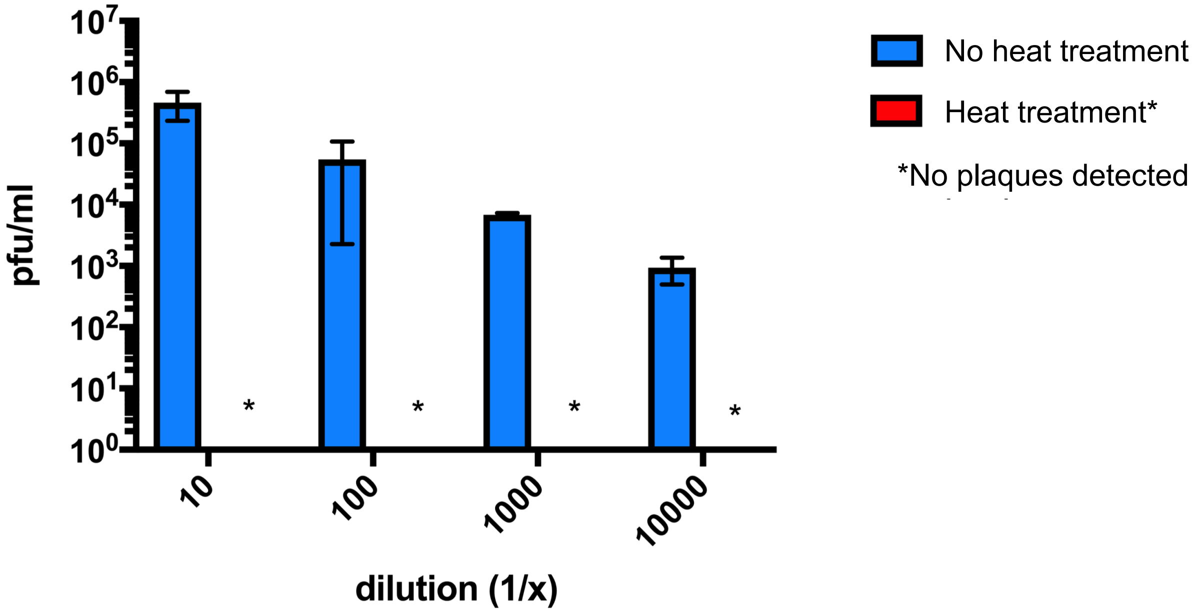


**Figure S1.** RVFV MP12 strain levels in heat treated and non-heat-treated milk. RVFV dilutions in milk were heated to 72 °C or kept at room temperature (negative control) for 15 min. Viral titres in the samples were determined by plaque assay. The data represent results of a representative experiment performed in triplicate.
